# Noise-processing by signaling networks

**DOI:** 10.1101/075366

**Authors:** Styliani Kontogeorgaki, Rubén J. Sánchez-García, Rob M. Ewing, Konstantinos C. Zygalakis, Ben D. MacArthur

**Affiliations:** Mathematical Sciences, University of Southampton, SO17 1BJ, UK; Biological Sciences, University of Southampton, SO17 1BJ, UK; School of Mathematics, University of Edinburgh, EH9 3FD, UK; Institute for Developmental Sciences, University of Southampton, SO17 1BJ, UK; Institute for Life Sciences, University of Southampton, SO17 1BJ, UK

## Abstract

Signaling networks mediate environmental information to the cell nucleus. To perform this task effectively they must be able to integrate multiple stimuli and distinguish persistent signals from transient environmental fluctuations. However, the ways in which signaling networks process environmental noise are not well understood. Here we outline a mathematical framework that relates a network’s structure to its capacity to process noise, and use this framework to dissect the noise-processing ability of signaling networks. We find that complex networks that are dense in directed paths are poor noise processors, while those that are sparse and strongly directional process noise well. These results suggest that while cross-talk between signaling pathways may increase the ability of signaling networks to integrate multiple stimuli, too much cross-talk may compromise the ability of the network to distinguish signal from noise. To illustrate these general results we consider the structure of the signaling network that maintains pluripotency in mouse embryonic stem cells, and find an incoherent feedforward loop structure involving Stat3, Tfcp2l1, Esrrb, Klf2 and Klf4 is particularly important for noise-processing. Taken together these results suggest that noise-processing is an important function of signaling networks and they may be structured in part to optimize this task.

## Introduction

Cellular identities are regulated by a variety of complex, interconnected, molecular regulatory networks, including signaling networks, metabolic networks and core transcriptional regulatory networks^1–9^. Signaling networks are of particular importance in maintaining robust cellular identities since they mediate noisy environmental information from the local cellular micro-environment to the cell nucleus^7, 10–14^. In order to perform this task effectively they must be able to transmit complex environmental information robustly, and failure to do this has been linked to cancer initiation and progression, as well as defects in embryonic development^15–17^. Much of what is known about signaling networks comes from the detailed reductionist analysis of their constituent signaling pathways. Several of these have been studied in great detail, and the core components and biochemical mechanisms of signal transduction in pathways such as Wnt, TGF-*β* and MAP Kinase signaling are now well defined. These signaling pathways function in a wide diversity of different biological processes and systems, and they are known to have a central role in maintaining pluripotency and specifying cell identities, for example^18, 19^. A long-standing question of interest is why, despite the myriad of biological processes and systems that involve signaling, there are only a few distinct, but widely conserved and re-used pathways^20^. An emerging feature that may in part explain this observation is that signaling pathway ‘modules’ are interconnected in many different ways and cross-talk between pathways has been shown for most signaling pathways, with mechanisms ranging from transcriptional activation of pathway components through to direct interactions between proteins in different pathways^20^. In addition to cross-talk between pathways, specific feedback mechanisms allow for the homeostatic control of signaling activity. A well defined example in the Wnt signaling pathway is the transcriptional activation of the Dickkopf proteins which negatively regulate Wnt signaling by binding to Wnt receptors in response to Wnt activation^21^. However, while much is now known about the function of specific signaling pathways, very little known about how cross-talk between pathways affects information-processing. In the context of signal processing it has been suggested that promiscuity in the protein-protein interactions is a major source of intrinsic noise, and that signaling pathways have evolved features (for example receptor clustering) to better distinguish signal from noise^22^. Although some studies have considered the ways in which noise propagates through regulatory networks^23–26^, the general mechanisms by which signaling networks distinguish persistent environmental signals from the noise that is inherent to the cellular micro-environment are still not well understood. To address this problem, here we outline a general mathematical framework which relates a network’s structure to its capacity to process noise, and use this framework to dissect the noise-processing ability of signaling networks. As an illustrative example we examine the noise-processing ability of the network that maintains pluripotency in mouse embryonic stem cells.

Embryonic stem (ES) cells are found naturally in the pre-implantation embryo and are able to give rise to all embryonic lineages, a property known as pluripotency. The molecular basis of pluripotency has been extensively studied, and it is now known that activation of a small number of core transcription factors – including Oct3/4, Sox2, and Nanog along with other secondary factors such as Myc, Klf4 and Lin28 – is sufficient to maintain the pluripotent state^5, 6, 27–34^. Indeed, forced expression of combinations of these factors in somatic cells is sufficient to induce pluripotency *de novo*^27, 29, 35^. Although this central transcriptional circuit is self-sustaining when shielded from external stimulation^36^, it is known that a network of signaling pathways which process extra-cellular environmental are also essential both to maintenance of, and exit from, the pluripotent state^37^. Importantly, While the core transcriptional circuity is broadly similar in mouse and human pluripotent cells^32^, their dependency on external signaling is markedly different: mouse ES cells are dependent on Lif/Stat signaling^38, 39^, Bmp^40^ and canonical Wnt^36^ to promote self-renewal, while Fgf/Erk signaling disrupts pluripotency^36, 41–43^; by contrast human ES cell self-renewal is independent of Lif^44^, yet requires requires Activin and Fgf^45, 46^ signaling and human ES cells undergo differentiation when exposed to Bmp^46^.

The remainder of the paper is organized as follows: we begin by outlining our general mathematical theory, as well as setting our assumptions, before establishing a mathematical formula that makes the connection between network structure and noise-processing explicit. To illustrate these results we then use this expression to investigate the structure of the regulatory network for pluripotency in mouse ES cells. This network is chosen since it is particularly well characterized^33^ and so constitutes a good test model. We find that certain elements in this network, particularly incoherent feedforward structures, are particularly important for its noise-processing ability. Interestingly, these elements are distinct from the core feedback structures that are known to maintain the pluripotent ground state^47^, suggesting that different portions of this network perform different regulatory tasks.

## Results

Our concern is with how a network *G* processes noise from an external source. In the context of signaling networks the nodes in the network are molecules in the signaling cascades, and edges are regulatory interactions (e.g. phosphorylation etc.) between molecules. Since signaling networks pass information from the cell exterior to the nucleus, we assume that the network *G* is inherently directed: the presence of an edge (*i, j*) indicates that node *i* exerts a regulatory effect on node *j* but not necessarily vice versa. Since regulatory interactions may be activatory or inhibitory we also allow each edge to have positive or negative weight representing the strength of activation or inhibition respectively. We denote the weight of edge (*i, j*) by *A_i j_*. Assuming that there are *n* nodes in *G* the *n×n* adjacency matrix ***A*** then describes the strength of all interactions in the system.

In general the regulatory interactions between nodes may be highly nonlinear. However, to better understand the relationship between network structure and function we will assume here that that the dynamics are linear. By doing so we are effectively considering the linearization near to a fixed point in the nonlinear dynamics; this rationale for studying the linear case has been taken elsewhere^48^. In the absence of external fluctuations, the dynamics of the system are described by the following system of ordinary differential equations (ODEs):

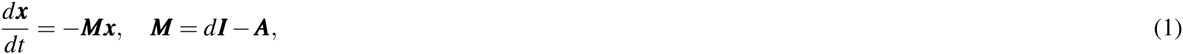

where ***I*** is the *n × n* identity matrix, and we have assumed that all nodes decay at the same rate *d*, which then sets a timescale for the dynamics. Without loss of generality we may take *d* = 1 since this may always be achieved by suitable re-scaling. Given the linearity of this system there are only two possible types of long-term behavior: convergence toward a stable fixed point or divergence to infinity. We will assume that only the first behavior can happen, *i.e.* convergence to a stable fixed point is the only physically realistic scenario. This occurs whenever the real parts of the eigenvalues of ***M*** are all strictly positive. Properties of the network *G* for which Eq. (1) admits a stable solution have been discussed at length, and it is known that sparse modular networks confer stability, for example^48, 49^. However, our concern here is not with stability *per se* but rather with the effect that external noise has on the magnitude of fluctuations around a stable equilibrium. To investigate this we will consider the following stochastic differential equation associated with Eq. (1)

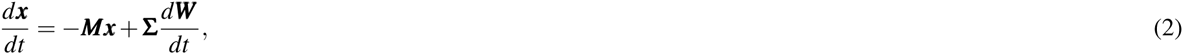

where ***W*** (*t*) is a standard *n*-dimensional Brownian motion. Eq. (2) describes a multivariate Ornstein-Uhlenbeck process. Whenever Eq. (1) admits a stable solution, Eq. (2) is ergodic and therefore admits a unique invariant measure^50^. Furthermore, by the linearity of Eq. (2) we know that ***x***(*t*) is distributed according to a multivariate normal with mean *e^—**M**t^ **x***(0) and covariance matrix ***K***(*t*) given by,

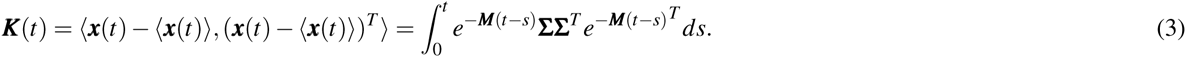

If Eq. (2) describes an ergodic process then

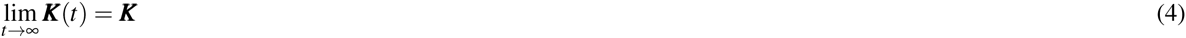

where ***K*** satisfies the Liapunov equation^50^

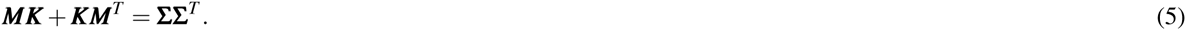

Although this is the standard formulation^50^, instead of working with Eq. (5) we will work directly with the Eq. (3) as it ultimately allows a more transparent assessment the effects of network structure on the stationary covariance of the system.

Since our purpose is to determine the way in which input noise is processed by the network *G* it is natural to consider a single noisy input to the system, which represents the fluctuating extra-cellular environment, and a single output, representing the computational core of the network. To do so we may, without loss of generality, chose a labeling of the nodes such that the first node is the noisy input and the *n*-th node is the output. Thus, we set **Σ** = (*σ*, 0, 0)^*T*^ and we are interested in calculating the variance of the *n*-th node in the network, which is given by *K_nn_*, relative to the magnitude of the input noise. If the input fluctuations carry no information, then this is a measure of the extent to which the network suppresses or amplifies random environmental fluctuations. If the fluctuations contain important environmental information, then this is a measure of the extent to which the target node can ‘sense’ this extra-cellular information. From Eq. (3) the limiting covariance, in index notation and using Einstein summation notation, is

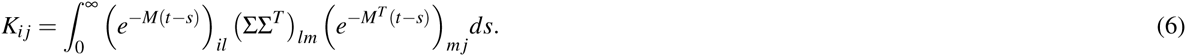

Now, since (ΣΣ^*T*^)_*lm*_ = *σ* ^2^ when *l* = *m* = 1 and zero otherwise, we have

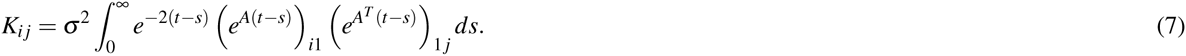

Although it is not immediately transparent, this expression connects strongly to the structure of the network via the fact that (*i, j*)-th entry of the exponential of the adjacency matrix of a network is a weighted sum of all walks between nodes *i* and *j*, and so is a simple measure of network ‘communicability’^51^. The weight *w*(*P*) of a walk *P* from node *i* to node *j* is the product of its edge weights, 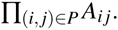 If *Pk* is the set of walks of length *k* between nodes *i* and *j* then the total weight of all walks from node *i* to node *j* with length 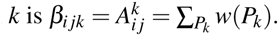 Using this notation, we may re-write the exponential terms in Eq. (7) as

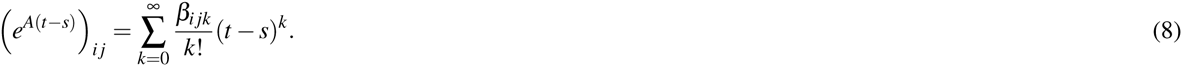

Making use of this connection and using the shorthand *β_ik_* = *β*_1*ik*_ we then obtain,

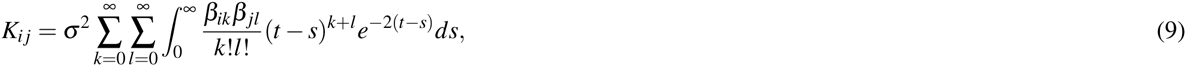

where we have adopted the convention that *β_i j_*_0_ = *δ_i j_*. Since

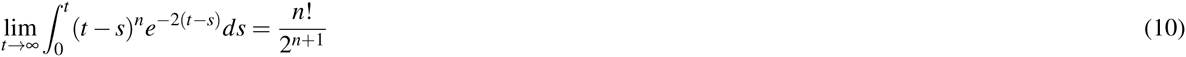

we may furthermore simplify Eq. (9) to

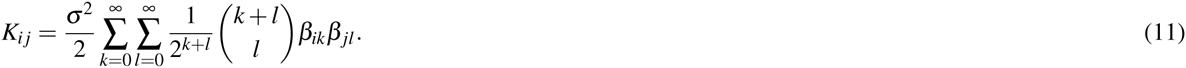

Since the variance of the noisy input is given by *K*_11_ = *σ* ^2^*/*2, we may investigate the noise-processing ability of the network by considering the ratio

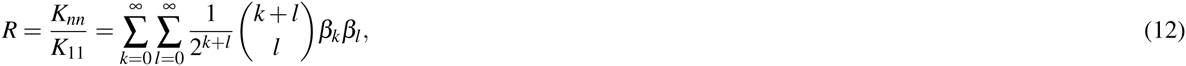

where we have further simplified notation by setting *β_nk_* = *β_k_*. When *R >* 1 noise is amplified by the network; when *R <* 1 noise is suppressed by the network. Importantly, if the process described by Eq. (2) is ergodic then *R* is finite and depends only on the structure of the network *G*. This formula therefore provides an explicit connection between network architecture and noise-processing; our interest is to determine how *R* is affected by different network architectures. To do so we note that Eq. (12) has a natural interpretation in terms of random walks on *G*, as follows.

Since each walk *P* from the input to the target has an associated weight *w*(*P*), a pair of (possibly intersecting) walks *P, Q* from the input to the target also has an associated weight *w*(*P, Q*) = *w*(*P*)*w*(*Q*), the product of the edge weights involved. If we write *P_k_* and *P_l_* for arbitrary walks from the input to the target of length *k*, and *l* respectively, then the product *β_k_β_l_* can be written as

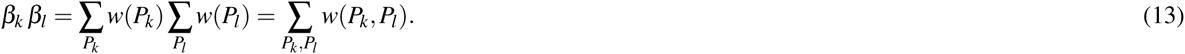

Substituting this into Eq. (12) and rewriting the second sum in terms of *m* = *k* + *l* gives,

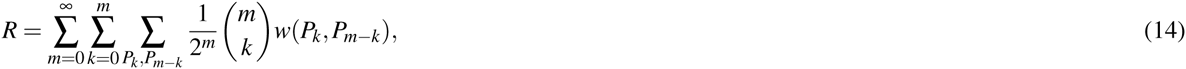

from which it can be seen that *R* is a weighted sum of all pairs of walks through *G* from the source node to the target, with the relative importance of each walk-pair determined by a coefficient drawn from a binomial distribution *B*(*m,* 1*/*2), where *m* is the length of the walk-pair. The appearance of binomial probabilities arises as the natural probability measure for pairs of random walkers on the network *G*. To see this consider two independent random walkers starting at the same time at the input node. At each time step, one walker is chosen with probability 1/2, and that walker moves through *G* choosing available edges with equal probability (*i.e.* if the walker is at node *i* then each outgoing edge from node *i* is chosen with probability 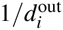 where 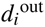 is the out-degree of node *i*). The probability that after precisely *m* = *k* + *l i* steps both walkers are at the target node is 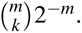. Thus, the inner sum in Eq. (14) is the expected weight of a pair of walks from input to target, with respect to the probability measure generated by two independent random walkers [the presence of two random walkers rather than one, as might be expected, arises from the fact that ***K*** depends upon two exponential terms, 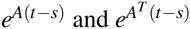, in Eq. (7)]. If *G* is a directed acyclic graph (in which there are no feedback loops), then all random walks have finite length and the the first sum in Eq. (14) has finitely many terms. However, if cycles are present in the network then random walks may be arbitrarily long and the first sum will correspondingly have infinitely many terms. Positive feedback loops add infinitely many positive terms to the sum, and therefore always serve to amplify noise with respect to similarly structured acyclic networks; negative feedback loops add both positive and negative terms to the sum and may amplify or reduce noise with respect to similarly structured acyclic networks, depending on the particular arrangement of inhibitory edges in the network.

It is worth noting here that although random walkers are a natural way to explore directed networks^52, 53^, if the matrix ***A*** is normal (that is, if it commutes with its transpose ***A**^T^*; a strong condition that is not typically satisfied by directed networks but does occur if the network *G* is undirected, for example), then a much simpler related result for the trace of the covariance matrix Tr(***K***) may be obtained. First, let us take the sum of the ratios *K_j j_/K*_11_ in Eq. (12) for all output nodes *j*,

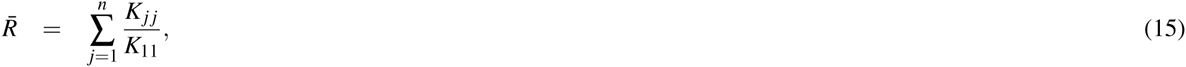

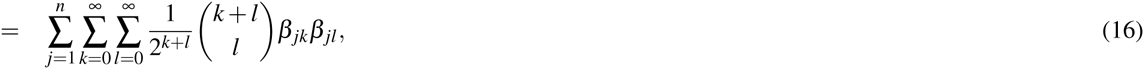

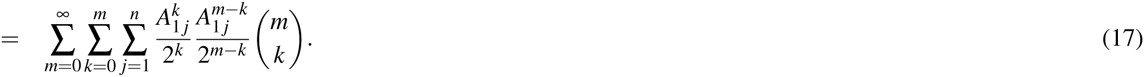

In general this sum cannot be simplified and we resort to interpreting in terms of random walkers, as above. However, if the matrices ***A*** and ***A**^T^* commute then we can use the binomial formula to expand the *m*-th power of the symmetric part of ***A*** as,

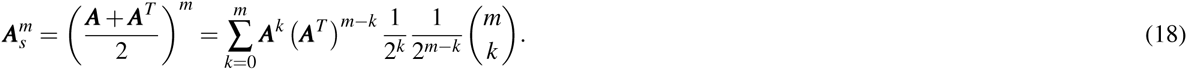

The (1, 1)-entry of the matrix above is precisely the double inner sum in Eq. (17), so we may write

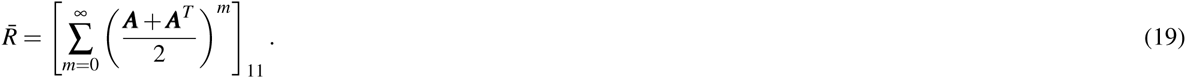

**Figure 1.**
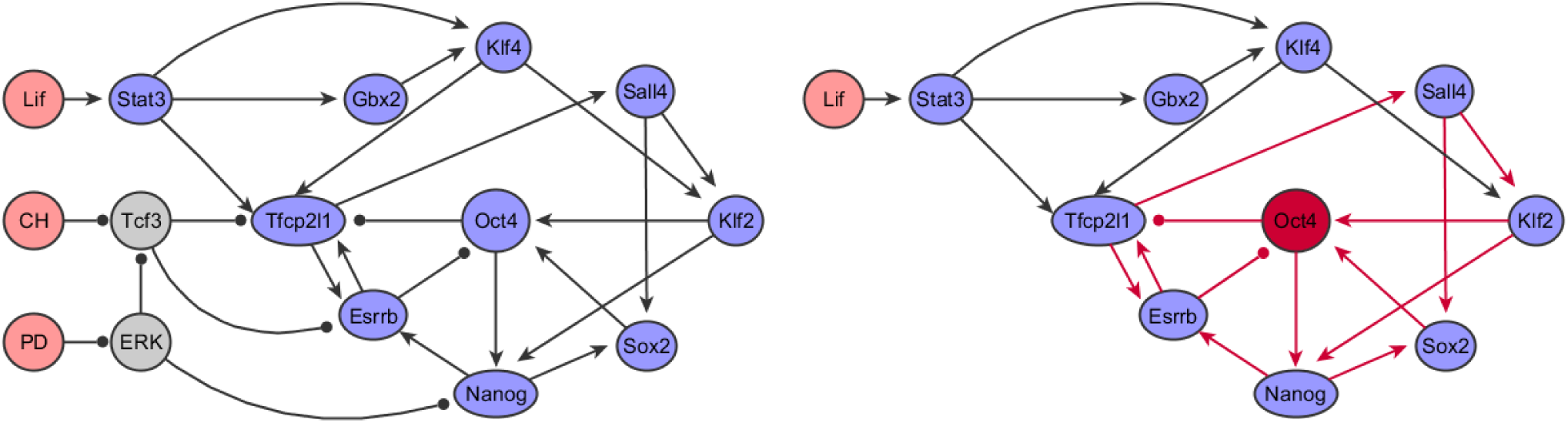
(Left) The regulatory network for pluripotency derived by Dunn and co-workers in^33^. (Right) The reduced network that we study here in which Lif is taken as a noisy input and Oct4 is taken as the target. Since CH and PD cannot be reached from Lif via a walk on this network, we can exclude these nodes, along with Tcf3 and ERK, from our analysis.

If 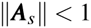 [a condition which implies stability in Eq. (1)], then the geometric series in Eq. (19) converges to

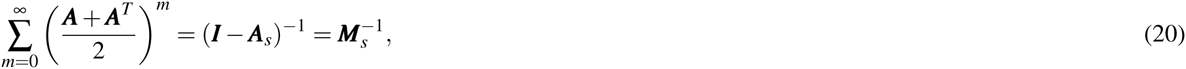

where ***M**_s_* = (***M*** + ***M***^*T*^)/2 is the symmetric part of ***M***. By taking into account all possible walks through the network, this is a simple variation on the exponential of the adjacency matrix as a measure of network communicability, although in this case the communicability of *G* is taken with respect to the analytic function (1 − *x*)^*−*1^, rather than exp(*x*)^54^. Using this result we finally obtain,

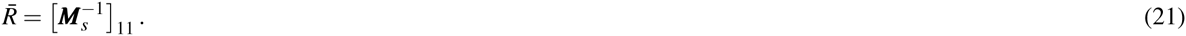

Thus, in order to determine the noise-processing ability of the network using this measure, we need only calculate the matrix inverse of the symmetric part of ***M***, and take the (1, 1)-entry (where without loss of generality the first node is the input node). This is result is strongly related to the Laplacian matrix of the network *G*: If ***A*** is symmetric and normalized so that each row (or column) sums to 1 then ***M***_*s*_ is precisely the normalized Laplacian of *G*, which is well-known to be closely related to network connectivity^55, 56^.

To illustrate how Eq. (12) works in practice, we now consider a couple of examples.

### Signaling cascade

The simplest example network is a signaling cascade, consisting of a chain of *m* = *n* − 1 interactions between *n* nodes. To facilitate a transparent illustration we shall assume that all the weights in the network are the same. In this case, we may index the nodes such that *A*_*i* *j*_ = *a* for edges (*i, i* + 1), where *i* = 1, 2*, …, n −* 1, and is zero otherwise. Eq. (12) then gives

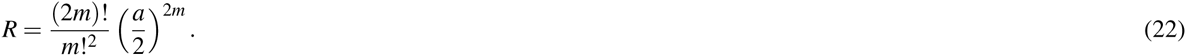

This result coincides with the expression derived in^57^. For large *m* we may use Stirling’s approximation to obtain

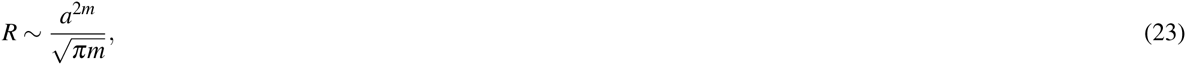

from which it can be seen that the signaling pathway suppresses noise if

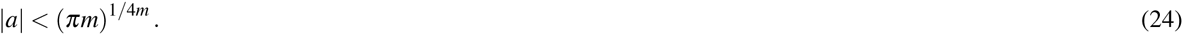

Three conclusions are apparent from this result: (1) since the variance of the target node depends upon the magnitude of the interaction strength squared, the sign of the interactions (*i.e.* whether they are activating or inhibiting) does not affect the ability of the cascade to process noise; (2) since *R* is monotonic decreasing with *m*, the effect of input noise diminishes with longer cascades (for fixed *a*); (3) if *a <* 1 then *R <* 1 for any *m* and noise is diminished by the cascade; while if *a >* 1 noise may be amplified by the cascade depending on the magnitude of *a* relative to *m*. In general since *R* decreases with *a* and *m*, this analysis suggests that long signaling cascades with weak interactions process noise better than short cascades with stronger interactions.

**Table 1:**
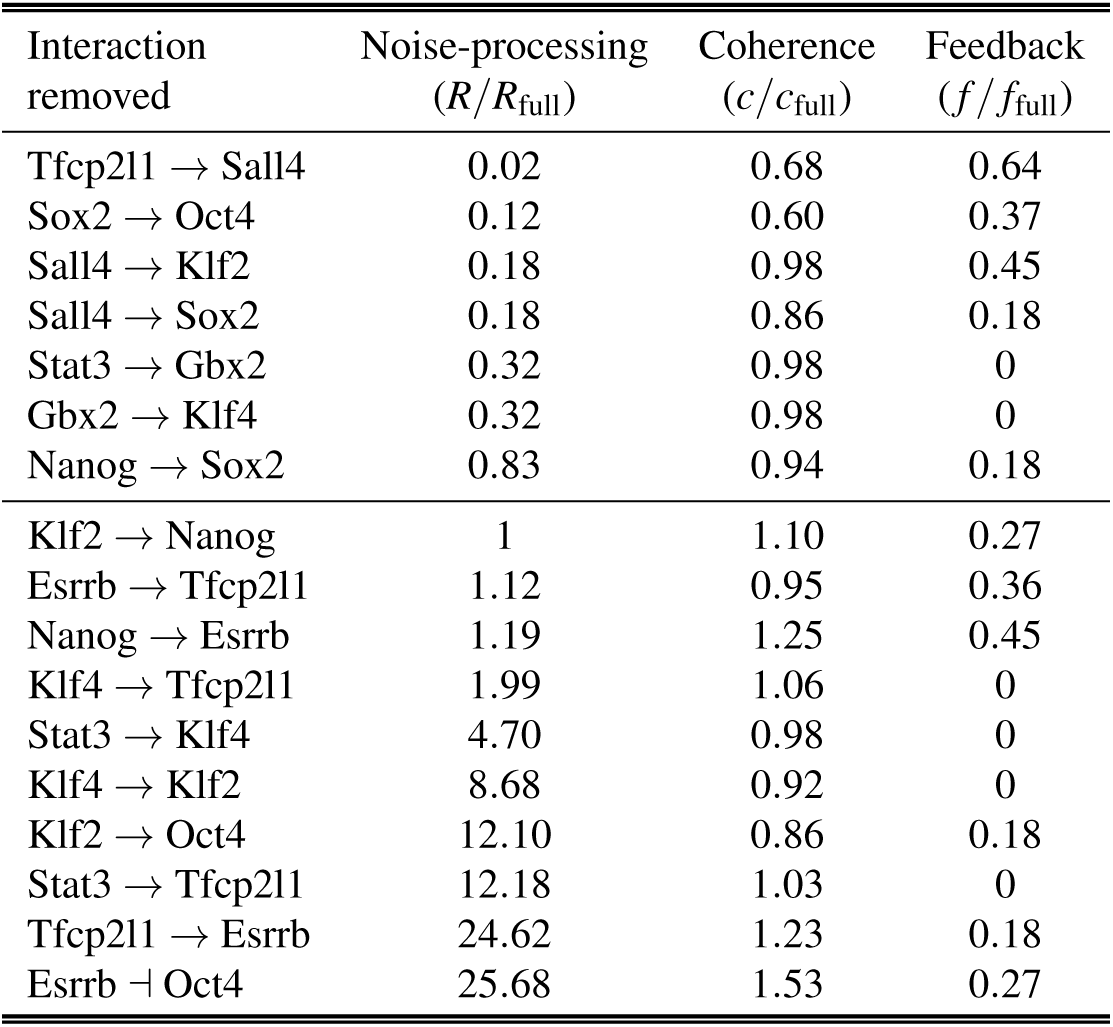
The effect that targeted removal of interactions on network noise-processing. The first column identifies the edge removed from the network; the second column shows the effect of targeted removal of the given edge on the ratio *R* by comparison with that of the unperturbed network; the third column shows the effect of targeted removal of the given edge has on network coherence; the fourth column shows the effect of targeted removal of the given edge has on network feedback. Edges that emanate from Oct4 do not contribute to the noise processing capacity of the network and their removal does not affect *R* so they are excluded from this table. Since all paths from Lif to Oct4 pass through the edge Lif → Stat3 its removal disconnects the network; this edge is also accordingly excluded from the table. Interactions are ordered by column 1.

### Feedforward loop

In reality signaling cascades do not operate in isolation; rather cross-talk between pathways means that many different paths from the input to the output may exist, and each may process different aspects of the extra-cellular signal. The simplest example of such a network is the feedforward loop motif, a commonly occurring structure in biological regulatory networks which is known to be involved with in a range of biological functions, including distinguishing persistent signals from noise^58^. The simplest feedforward loop consists of three nodes with two paths from the source (node 1) to the target (node 3): one direct and one indirect, via an intermediary (node 2). In this case, assuming that all interactions are of equal weight we obtain

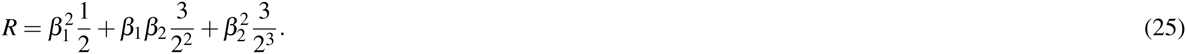

By contrast to the simple signaling cascade, the sign of the edges in the feedforward loop do affect its noise-processing ability. If all edges are positive then *β*_1_, *β*_2_ *>* 0 (all paths in the network are positive) and the target receives a consistent signal from the source. In this case the feedforward loop is said to be coherent^58^. However, if *β*_1_ *<* 0 or *β*_2_ *<* 0 (which occurs if either one or three of the edges is negative) then the target receives a inconsistent signal from the source. In this case, the feedforward loop is said to be incoherent^58^. Denoting the noise-processing ratios in the coherent and incoherent cases by *R*^+^ and *R^−^* respectively, it follows from Eq. (25) that *R*^+^ *> R^−^*, and therefore that the incoherent feedback loop is better at processing noise. This general conclusion typically holds for more complex feedforward structures which may contain a larger number of longer paths from the source to the target: incoherence of paths through the network tends to lead to better noise-processing.

### Noise-processing by stem cells

In order to apply these general results we now consider signal transduction in the regulatory network for pluripotency. The skeleton of this network has recently been inferred from analysis of correlations between expression patters of important regulatory factors^33^ and is illustrated in Fig. 1. The behavior of this network is determined by input from three extra-cellular factors commonly added to ES cell culture media preparations: the cytokine leukemia inhibitory factor (Lif), and selective inhibitors of glycogen synthase kinase 3 (TGF-*β*) (Chiron99021, denoted CH) and mitogen-activated protein kinase kinase (Mek) (PD0325901, denoted PD). Since it is known that stimulation of Lif signaling is sufficient to maintain pluripotency *in vitro*, we will chose Lif as the noisy source in our analysis. When Lif is present in the extra-cellular environment it binds to the Stat3 receptor^59, 60^, and activates signaling pathways that stimulate the core transcriptional regulatory network for pluripotency in mouse ES cells^34, 61^. At the center of this core network are the trio of transcription factors, Oct4, Sox2 and Nanog^28, 62, 63^. Since Oct4 is well-established as the most central factor in this core, we will consider the propagation of a noisy signal from (the input) Lif through the network to (the output) Oct4. Although the sign of the interactions (i.e. whether they are activatory or inhibitory) is known^33^ their strength is unknown. In the absence of this information we assume that all interactions are of equal unit strength since this represents the most economic model. To investigate how network structure affects noise-processing, we sought to determine the how the ratio *R* changes upon targeted removal of different interactions from this network by comparing the *R* values of perturbed networks with that of the unperturbed network (denoted *R*_full_). To uncover the structural determinants of noise-processing we also calculated two simple network measures based upon our interpretation of Eq. (12): (1) *c* = *p*^+^ (*p*^+^ + *p*^*−*^), where *p*^+^ and *p*^−^ are the number of positive and negative paths from Lif to Oct4 respectively. This is a simple measure of structural coherence; and (2) *f*, the total number of feedback loops in the network, as a measure of network complexity. To determine how network properties varied with removal of specific edges, we also determined how these measures changed upon targeted removal of different interactions from the network by comparing values with those of the unperturbed network (denoted *c*_full_ and *f*_full_ respectively). The results of analysis are summarized in Table 1.

**Figure 2.**
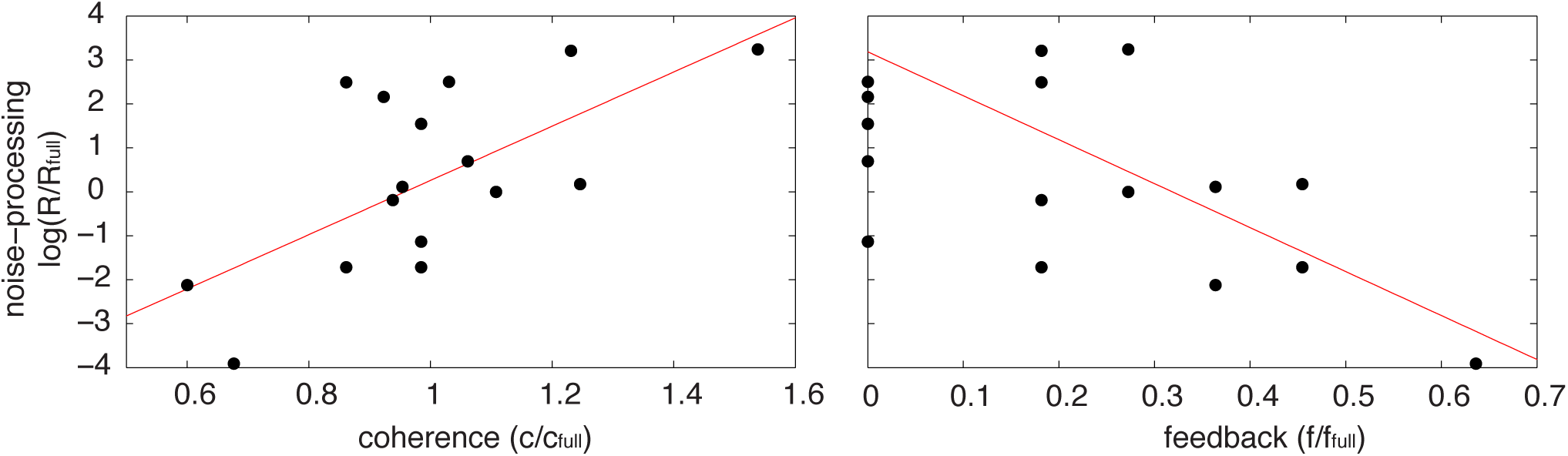
Plots of the data from Table 1. Removal of edges that result in an increase of coherence in the network tend to diminish the system’s noise-processing ability, while removal of edges which reduce the overall feedback structure of the network tend to improve the system’s noise-processing ability. Red lines show linear regression.

The shortest paths from Lif to Oct4 in this network have length 4; there are two such paths: (1) Lif → Stat3 → Tfcp2l1 → Esrrb ⊣ Oct4; and (2) Lif → Stat3 → Klf4 → Klf2 → Oct4. The first of these paths in negative (due to the inhibitory interaction Esrrb ⊣ Oct4) while the second is positive. Thus, when taken together this pair of paths forms an incoherent feedforward loop; since these are the shortest paths in the network, we anticipate from Eq. (12) that this incoherent feedforward loop will have an important role in noise-processing in this network. Indeed, this is what is observed: if any of the elements of this structure are removed, then the noise-processing capacity of this network is severely inhibited and the ratio *R* increases substantially (see Table 1, Fig. 2). By contrast, we also anticipate that since feedback loops introduce arbitrarily long walks in the network [and therefore contribute infinitely many terms to the sum in Eq. (12)] removal of edges which participate in the feedback structure of the network will result in a substantial reduction in its noise-processing capacity. Again, this is what is observed: when edges which participate in large numbers of feedback loops are removed, the ratio *R* decreases substantially (see Table 1, Fig. 2). Taken together, these results indicate that those interactions in the feedback rich core transcriptional circuitry that are needed to maintain a self-perpetuating pluripotent identity^5, 47^ tend to amplify extrinsic noise. To compensate, a distinct set of interactions between auxiliary factors structured into a set of incoherent feedforward loops suppress environmental noise, and ensure that environmental signals are robustly mediated to this core circuit.

## Discussion

In this paper we have derived a simple expression that relates a network’s structure to its noise-processing ability. This expression is easily calculated even for large networks [particularly if the network is undirected, see Eq. (21)], so provides an economic measure that may be used to examine the structure of naturally occurring networks, and guide the design of man-made networks. To illustrate these results we have considered the structure of the network that maintains pluripotency in mouse ES cells, and found that important network structures, distinct from those that maintain the core pluripotent state, are responsible for noise processing in this system, suggesting that different features of this network are responsible for different regulatory tasks. Accordingly, we anticipate that the structure of many natural networks may be determined, in part, to optimally process noise. It will be interesting to elucidate the extent to which cross-talk between pathways in natural networks, which typically have to process multiple complex signals, is shaped by the trade-off between signal integration and noise-processing.

## Acknowledgments

This work was funded by BBSRC Grant No. BB/L000512/1 and by PhD funding from the Institute for Life Sciences, University of Southampton. The authors would like to thank Enrico Mossotto, Rosanna Smith and Patrick Stumpf for useful feedback on the manuscript.

## Author contributions statement

SK, RSG, KCZ and BDM conducted the mathematical modeling and network analysis. SK, RSG, RE, KCZ and BDM interpreted the results. All authors wrote the paper.

## References

1. Barabási, A. L. & Oltvai, Z. N. Network biology: Understanding the cell’s functional organization. Nat. Rev. Genet. 5, 101–113 (2004).

2. Tkačik, G. & Bialek, W. Cell biology: Networks, regulation, pathways. In Encyclopedia of complexity and systems science, 719–741 (Berlin: Springer, 2009).

3. Newman, M. E. J. Networks. An introduction (OUP, 2010).

4. Estrada, E. The structure of complex networks (OUP, 2011).

5. Macarthur, B. D., Ma’ayan, A. & Lemischka, I. R. Systems biology of stem cell fate and cellular reprogramming. Nat. Rev. Mol. Cell Biol. 10, 672–681 (2009).

6. Martello, G. & Smith, A. The nature of embryonic stem cells. Annu. Rev. Cell Dev. Biol. 30, 647–675 (2014).

7. Jeong, H., Mason, S. P., Barabási, A. L. & Oltvai, Z. N. Lethality and centrality in protein networks. Nature 411, 41–42 (2001).

8. Blais, A. & Dynlacht, B. D. Constructing transcriptional regulatory networks. Genes Dev. 19, 1499–1511 (2005).

9. Alberts, B. et al. Molecular biology of the cell (Garland science, Taylor and Francis Group, LLC, 2002).

10. Adler, E. M., Gough, N. R. & Ray, L. B. 2015:) Signaling breakthroughs of the year. Sci. Signal. 9, eg1 (2016).

11. Irish, J. M. et al. Single cell profiling of potentiated phospho-protein networks in cancer cells. Cell 118, 217–228 (2015).

12. Verkaar, F., Cadigan, K. M. & Amerongen, V. R. Celebrating 30 years of wnt signaling. Sci. Signal. 5, mr2 (2012).

13. Levine, E. & Hwa, T. Stochastic fluctuations in metabolic pathways. Proc. Natl. Acad. Sci. USA. 104, 9224–9229 (2007).

14. Schreiber, S. L. & Bernstein, B. E. Signaling network model of chromatin. Cell 111, 771–778 (2015).

15. Song, J. et al.A protein interaction between β-catenin and dnmt1 regulates wnt signaling and dna methylation in colorectal cancer cells. Mol. Cancer Res. 13, 969–981 (2015).

16. Valenta, T., Hausmann, G. & Basler, K. The many faces and functions of β-catenin. EMBO J. 31, 2714–36 (2012).

17. Clevers, H., Loh, K. M. & Nusse, R. An integral program for tissue renewal and regeneration: Wnt signaling and stem cell control. Science 346, 1248012 (2014).

18. Niwa, H. Wnt: What’s needed to maintain pluripotency? Nat. Cell Biol. 13, 1024–1026 (2011).

19. Kim, H. et al. Modulation of β-catenin function maintains mouse epiblast stem cell and human embryonic stem cell self-renewal. Nat. Commun. 4, 2403 (2013).

20. Attisano, L. & Wrana, J. L. Signal integration in tgf-β, wnt, and hippo pathways. F1000Prime Rep. 5 (2013).

21. Niida, A. et al. Dkk1, a negative regulator of wnt signaling, is a target of the beta-catenin/tcf pathway. Oncogene 23, 8520–8526 (2004).

22. Ladbury, J. E. & Arold, S. T. Noise in cellular signaling pathways: causes and effects. Trends Biochem. Sci. 37, 173–178 (2012).

23. Pedraza, J. M. & van Oudenaarden, A. Noise propagation in gene networks. Science 307, 1965–1969 (2005).

24. Paulsson, J. Summing up the noise in gene networks. Nature 427, 415–418 (2004).

25. Hornung, G. & Barkai, N. Noise propagation and signaling sensitivity in biological networks: A role for positive feedback. PLoS Comput. Biol. 4, e8 (2008).

26. Hooshangi, S. & Weiss, R. The effect of negative feedback on noise propagation in transcriptional gene networks. Chaos 16, 026108 (2006).

27. Takahashi, K. & Yamanaka, S. Induction of pluripotent stem cells from mouse embryonic and adult fibroblast cultures by defined factors. Cell 126, 663–676 (2006).

28. Boyer, L. A. et al. Core transcriptional regulatory circuitry in human embryonic stem cells. Cell 122, 947–956 (2005).

29. Yu, J. et al. Induced pluripotent stem cell lines derived from human somatic cells. Science 318, 1917–1920 (2007).

30. Dahéron, L. et al. LIF/STAT3 signaling fails to maintain self-renewal of human embryonic stem cells. Stem Cells 22, 770–778 (2004).

31. Ye, S., Li, P., Tong, C. & Ying, Q. Embryonic stem cell self-renewal pathways converge on the transcription factor Tfcp2l1. EMBO J. 32, 2548–60 (2013).

32. Kim, J., Chu, J., Shen, X., Wang, J. & Orkin, S. H. An extended transcriptional network for pluripotency of embryonic stem cells. Cell 132, 1049–1061 (2008).

33. Dunn, S. J., Martello, G., Yordanov, B., Emmott, S. & Smith, A. G. Defining an essential transcription factor program for naïve pluripotency. Science 344, 1156–60 (2014).

34. Huang, G., Ye, S., Zhou, X., Liu, D. & Ying, Q. L. Molecular basis of embryonic stem cell self-renewal: From signaling pathways to pluripotency network. Cell. Mol. Life Sci. 72, 1741–1757 (2015).

35. Takahashi, K. et al. Induction of pluripotent stem cells from adult human fibroblasts by defined factors. Cell 131, 861–872 (2007).

36. Ying, Q.-L. et al. The ground state of embryonic stem cell self-renewal. Nature 453, 519–523 (2008).

37. Stumpf, P. S., Ewing, R. & MacArthur, B. D. Single-cell pluripotency regulatory networks. Proteomics 16, 2303–2312 (2016).

38. Williams, R. L., Hilton, D. J. & Nicolai, N. A. Myeloid leukaemia inhibitory factor maintains the developmental potential of embryonic stem cells. Nature 336, 15 (1988).

39. Smith, A. G. et al. Inhibition of pluripotential embryonic stem cell differentiation by purified polypeptides. Nature 336, 688–690 (1988).

40. Ying, Q.-L., Nichols, J., Chambers, I. & Smith, A. Bmp induction of id proteins suppresses differentiation and sustains embryonic stem cell self-renewal in collaboration with stat3. Cell 115, 281–292 (2003).

41. Burdon, T., Stracey, C., Chambers, I., Nichols, J. & Smith, A. Suppression of shp-2 and erk signalling promotes self-renewal of mouse embryonic stem cells. Dev. Biol. 210, 30–43 (1999).

42. Stavridis, M. P., Lunn, J. S., Collins, B. J. & Storey, K. G. A discrete period of fgf-induced erk1/2 signalling is required for vertebrate neural specification. Development 134, 2889–2894 (2007).

43. Kunath, T. et al. Fgf stimulation of the erk1/2 signalling cascade triggers transition of pluripotent embryonic stem cells from self-renewal to lineage commitment. Development 134, 2895–2902 (2007).

44. Sato, N., Meijer, L., Skaltsounis, L., Greengard, P. & Brivanlou, A. H. Maintenance of pluripotency in human and mouse embryonic stem cells through activation of wnt signaling by a pharmacological gsk-3-specific inhibitor. Nat. Med. 10, 55–63 (2004).

45. Amit, M. et al. Clonally derived human embryonic stem cell lines maintain pluripotency and proliferative potential for prolonged periods of culture. Dev. Biol. 227, 271–278 (2000).

46. Xu, R.-H. et al. Basic FGF and suppression of BMP signaling sustain undifferentiated proliferation of human es cells. Nat. Methods 2, 185–190 (2005).

47. MacArthur, B. D. et al. Nanog-dependent feedback loops regulate murine embryonic stem cell heterogeneity. Nat. Cell Biol. 14, 1139–1147 (2012).

48. May, R. M. Will a large complex system be stable? Nature 238, 413–414 (1972).

49. Allesina, S. & Tang, S. Stability criteria for complex ecosystems. Nature 483, 205–208 (2012).

50. Gardiner, C. W. Stochastic methods: a handbook for the natural and social sciences (Berlin: Springer, 2009).

51. Estrada, E. & Higham, D. J. Network properties revealed through matrix functions. SIAM Rev. 52, 696–714 (2010).

52. Rosvall, M. & Bergstrom, C. T. Maps of random walks on complex networks reveal community structure. Proc. Natl. Acad. Sci. USA. 105, 1118–1123 (2008).

53. Chung, F. Laplacians and the Cheeger inequality for directed graphs. Ann. Comb. 9, 1–19 (2005).

54. Estrada, E. & Higham, D. J. Network properties revealed through matrix functions. SIAM Rev. 52, 696–714 (2010).

55. Mohar, B. The Laplacian spectrum of graphs. In Graph theory, combinatorics, and applications., Wiley-Intersci. Publ., 871–898 (Wiley: New York, 1991).

56. Luxburg, U. V. A tutorial on spectral clustering. Stat. Comput. 17, 395–416 (2006).

57. Anderson, D. F., Mattingly, J. C., Nijhout, H. F. & Reed, M. C. Propagation of fluctuations in biochemical systems, i: Linear ssc networks. Bull. Math. Biol. 69, 1791–1813 (2007).

58. Alon, U. An introduction to systems biology: design principles of biological circuits (Chapman Hall, 2007).

59. Matsuda, T. et al. STAT3 activation is sufficient to maintain an undifferentiated state of mouse embryonic stem cells. EMBO J. 18, 4261–4269 (1999).

60. Niwa, H., Burdon, T., Chambers, I. & Smith, A. Self-renewal of pluripotent embryonic stem cells is mediated via activation of STAT3. Genes Dev. 12, 2048–2060 (1998).

61. Yu, J. & Thomson, J. Pluripotent stem cell lines. Genes Dev. 22, 1987–1997 (2008).

62. Niwa, H., Miyazaki, J. & Smith, A. G. Quantitative expression of Oct-3/4 defines differentiation, dedifferentiation or self-renewal of ES cells. Nat. Genet. 24, 372–376 (2000).

63. Nichols, J. et al. Formation of pluripotent stem cells in the mammalian embryo dependes on the POU transcription factor Oct4. Cell 95, 379–391 (1998).

